# Point-Of-Need One-Pot Multiplexed RT-LAMP Test For Detecting Three Common Respiratory Viruses In Saliva

**DOI:** 10.1101/2025.03.07.642108

**Authors:** Aneesh Kshirsagar, Dean DeRosa, Anthony J. Politza, Tianyi Liu, Ming Dong, Weihua Guan

## Abstract

Respiratory viral infections pose a significant global public health challenge, partly due to the difficulty in rapidly and accurately distinguishing between viruses with similar symptoms at the point of care, hindering timely and appropriate treatment and limiting effective infection control and prevention efforts. Here, we developed a multiplexed, non-invasive saliva-based, reverse transcription loop-mediated isothermal amplification (RT-LAMP) test that enables the simultaneous detection of three of the most common respiratory infections, severe acute respiratory syndrome coronavirus 2 (SARS-CoV-2), Influenza (Flu), and respiratory syncytial virus (RSV), in a single reaction via specific probes and monitored in real-time by a machine-learning-enabled compact analyzer. Our results demonstrate that the multiplexed assay can effectively detect three target RNAs with high accuracy. Further, testing with spiked saliva samples showed strong agreement with reverse transcription polymerase chain reaction (RT-PCR) assay, with area under the curve (AUC) values of 0.82, 0.93, and 0.96 for RSV, Influenza, and SARS-CoV-2, respectively. By enabling the rapid detection of respiratory infections from easily collected saliva samples at the point of care, the device presented here offers a practical and efficient tool for improving outcomes and helping prevent the spread of contagious diseases.

**Significance:** This research presents an innovative approach to respiratory infection diagnostics by combining a one-pot isothermal molecular test with machine learning-based analysis to simultaneously detect SARS-CoV-2, Influenza, and RSV in saliva samples. The battery-powered portable analyzer features novel machine-learning-assisted fluorescence detection for multiplexed reporter quantification, eliminating the need for traditional filter-based optical components and enabling adaptation to new targets without hardware changes. The test demonstrates high accuracy in detecting single and co-infections in spiked saliva samples, providing a rapid, cost-effective point-of-need solution. This tool can expand testing access, improve patient outcomes, and support more effective disease control, particularly in resource-limited or decentralized healthcare settings.

## Introduction

The accurate diagnosis of respiratory infections is particularly challenging when multiple pathogens share similar clinical symptoms. Seasonal circulations of Influenza A (IAV), Influenza B (IBV), and Respiratory Syncytial Virus (RSV) were the most well-known examples of such viral infections until the widespread emergence of the Severe Acute Respiratory Syndrome Coronavirus 2 (SARS-CoV-2) virus(1). These pathogens present similar or overlapping symptoms such as fever, cough, dyspnea, myalgia, and the typical changes in chest radiology images, including ground-glass opacities(2). As a result, healthcare providers may find it challenging to rely solely on syndromic diagnosis to pinpoint the exact causative pathogen, which may lead to delayed or purely symptomatic treatment(3). Inappropriate or insufficient treatment can lead to worsening symptoms or related complications, particularly in high-risk patient groups(4), resulting in longer recovery times and compromised outcomes(5).

In addition, delays in accurate diagnosis can hinder public health measures such as infection control or quarantine protocols, potentially allowing the further spread of highly contagious pathogens. This challenge was particularly evident during the 2022-2023 US ‘tripledemic’ when widespread cases of SARS-CoV-2, Influenza, and RSV coincided, overwhelming clinics and hospitals(6). Moreover, co-infection by two viruses has been reported worldwide(7–11). Identifying and managing co-infections is crucial, particularly for high-risk cases, as different pathogens may require distinct treatments: remdesivir(12), molnupiravir(13), and nirmatrelvir co-formulated with ritonavir(14) for COVID-19, oral oseltamivir, inhaled zanamivir, or intravenous peramivir for Influenza(7), and usually only symptomatic treatment for RSV(15). Thus, a reliable and timely molecular diagnosis of the causative agent(s) is essential to ensure that the most suitable treatment is administered within the appropriate window of 5–7 days for COVID-19 and 48 hours for Influenza after the onset of symptoms(13, 16).

Current diagnostic practices to differentiate between infections often involve respiratory panels(17–19) that rely primarily on parallel polymerase chain reactions (PCR)(20) or isothermal amplification techniques such as loop-mediated isothermal amplification (LAMP)(21), recombinase polymerase amplification (RPA)(22), helicase-dependent amplification (HDA)(23), or specific high-sensitivity enzymatic reporter unlocking (SHERLOCK)(24), which may be conducted as sequential single-plex(25–28) or multiplex assays(29–32). Currently, all these tests rely on nasal or nasopharyngeal swabs, which require careful sample collection from skilled practitioners and could be uncomfortable, leading to patient hesitance due to discomfort or risk of minor injury(33, 34). Additionally, errors during sample collection, such as insufficient genetic material, can lead to inaccurate or inconclusive results. During the COVID-19 pandemic, saliva emerged as a promising and simplified alternative to nasal/nasopharyngeal swabs for SARS-CoV-2 detection, offering comparable sensitivity even when stored without preservative reagents(35–39). Saliva collection is non-invasive, easily self-administered, and generally well-tolerated, which could enhance patient compliance.

Our previous work demonstrated a sample-to-answer test that uses saliva and reverse transcription loop-mediated isothermal amplification (RT-LAMP) to detect SARS-CoV-2 in a single-plex format with three parallel reaction chambers, which could be adapted for multiple targets(40). RT-LAMP often relies on non-specific detection methods such as intercalating dyes or colorimetric approaches, making multiplexing particularly challenging(41). Consequently, most studies detecting multiple targets rely on parallel reactions(42–44). While one-pot reactions offer advantages such as potentially improved sensitivity and eliminating the risk of non-uniform sample splitting, efforts to develop more specific detection strategies remain limited(41, 45–47). Notably, despite the growing demand for point-of-need tests, these strategies have not yet been widely implemented in such settings, which highlights a critical gap. Given RT-LAMP’s potential and the ease of using saliva as a sample type, a one-pot point-of-need saliva-based nucleic acid test (NAT) leveraging isothermal amplification could enable large-scale adoption, home testing, and widespread screening for respiratory infections.

In this work, we developed a highly specific and multiplexed RT-LAMP-based test that utilizes saliva for the simultaneous detection of infections such as SARS-CoV-2, Influenza (Flu), and RSV in a single reaction. The RT-LAMP assay uses the detection of amplification by release of quenching (DARQ) technique(45) to allow multiplexed detection via fluorescently labeled primers specifically for a target. Complementing this assay, we developed a compact, battery-powered instrument capable of detecting multiple fluorophores simultaneously, empowered by machine learning to estimate time-varying fluorophore concentrations. In contrast to traditional systems with excitation and emission filters, which may limit the number of detectable targets, our design does not rely on expensive optical filters for specificity and eliminates the need for hardware reconfiguration when incorporating new or additional fluorophores, allowing for easy future adaptation to new pathogens or variants. We demonstrate the assay’s potential to address the critical need for differentiating between respiratory infections and identifying co-infections by running the optimized multiplexed RT-LAMP assay on mock saliva samples spiked with single or combinations of target RNAs. This system can potentially improve diagnostic efficiency, enabling more precise identification of co-infections and pinpointing the causative pathogens in point-of-need settings, improving timely treatment decisions and patient outcomes.

## Results

### A saliva-based point-of-need diagnostic test for multiplexed respiratory tri-virus detection

To accurately identify the possible causative virus leading to respiratory symptoms at the point of need, we propose a diagnostic test, the workflow of which is given in **Figure 1a**. It involves the collection of saliva, extraction of RNA using the Qiagen RNA extraction mini kit performed on a custom-developed potable centrifuge, and finally, running the one-pot multiplexed RT-LAMP reaction on a portable nucleic acid testing device while using all the relevant primer sets and distinct fluorescent probes to distinguish between SARS-CoV-2, Flu, RSV or no infection.

**Figure 1.**
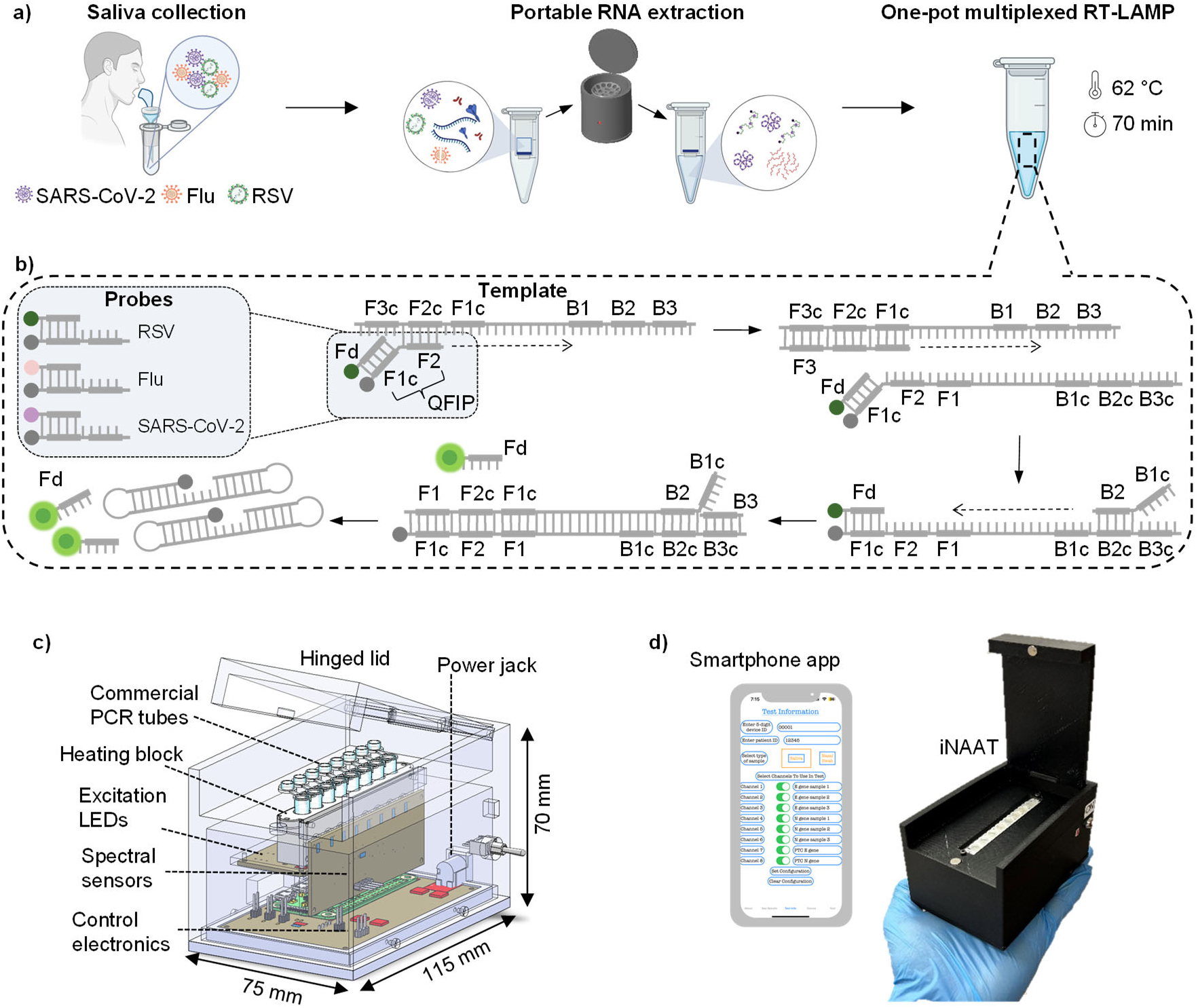
An overview of the test and the analyzer developed for multiplexed monitoring of RT-LAMP reaction. (a) The test workflow involves i) saliva collection, ii) portable RNA extraction, and iii) multiplexed RT-LAMP using specific primers and distinct fluorescent probes, enabling one-pot multiplexing. Multiplexing allows simultaneous detection and differentiation of RSV, Flu, SARS-CoV-2, or no infection within a single reaction. (b) Schematic representing the mechanism of DARQ RT-LAMP for multiplex detection using fluorescently labeled probes. A quencher-tagged Forward Internal Primer (QFIP) is annealed to a fluorescently labeled probe (Fd) complementary to the F1c region before adding to the reaction. The fluorescent probe remains quenched during initiation at the F2 region, and subsequent displacement of the quenched strand is triggered by the hybridization of the F3 primer, leading to fluorescence release as the Fd probe separates from QFIP when the Backward Internal Primer (BIP) initiates backward strand formation. Fluorescence production increases as more amplicons are generated during LAMP until a plateau is reached. (c) 3D CAD diagram of the developed analyzer, which can run up to eight multiplexed reactions in parallel and is compatible with traditional PCR tubes. (d) Pictorial representation of fabricated analyzer and smartphone running the custom- developed app.

To allow one-pot multiplexing, we make use of distinct fluorophores for each viral target, and **Figure 1b** schematically outlines the DARQ RT-LAMP mechanism, which allows multiplex detection via fluorescently labeled probes, initially described by Tanner et al.(45). Like traditional LAMP, DARQ LAMP consists of six primers that target eight regions of the template cDNA with the Forward Internal Primer (FIP) tagged with a quencher on the 5’ end (termed QFIP) and a corresponding probe complementary to only the F1c region and tagged with a fluorophore (termed Fd). Fd and QFIP are annealed together by heating to 95 °C and slowly cooling to room temperature at ∼1 °C/min in a controlled manner before adding to the amplification reaction. Like traditional LAMP, DARQ LAMP initiates at the F2c region while the fluorescent probe remains annealed and quenched. This strand is then displaced because of the hybridization of the F3 primer to initiate a new strand. Meanwhile, the previously displaced strand experiences hybridization at the B2c region, and upon extension, the annealed Fd is released from QFIP to produce fluorescence. As LAMP progresses, more amplicons are generated to produce more fluorescence until a plateau is reached. Please note that the reverse transcription step and other intermediate steps within LAMP are not shown for simplicity. The one-pot reaction is performed on a handheld analyzer, iNAAT, and **Figure 1c** presents its 3D render. The three main modules of the developed analyzer are detailed in **Supplementary Figure S1a**. The analyzer is designed to run eight reactions in parallel, is compatible with traditional PCR tubes, and has a power consumption of ∼5 Wh such that it can be powered either by a commercial rechargeable battery pack to allow up to 12 hrs. of in-field use, or by a wall adapter to allow potentially continuous lab use. The optical assembly used in this analyzer has been described in detail earlier(48); it consists of an RGB LED and an eight-channel spectral sensor mounted perpendicular to each other and using a deep learning model trained on a set of calibration data acquired from mixtures of chosen fluorophores, the analyzer can accurately predict the concentration of constituent fluorophores in a reaction as output. These predicted concentrations are shared with an iOS app over Bluetooth developed to provide test instructions, acquire data, and make positive and negative calls to interpret the test results. **Figure 1d** shows the in-house fabricated analyzer and a smartphone running the app, and **Supplementary Video V1** demonstrates the test workflow.

### DARQ RT-LAMP assays’ cross-reactivity validation

To develop a multiplexed RT-LAMP assay, we first verified that each of the premier sets targeting RSV, Flu, and SARS-CoV-2 are specific, i.e., do not detect any of the other targets and do not interact with each other to form primer-dimers that may lead to false amplification. **Figure 2a** shows the specific regions of the three RNA targets that are amplified and detected during the single-plex DARQ RT-LAMP reactions, N, HA, and N1, for RSV, Flu, and SARS-CoV-2 respectively. The primer sequences used in these assays have been previously reported(32, 49, 50), with modifications made to the FIP primer, incorporating a quencher at the 5’ end (QFIP), and designing a complementary fluorescence labeled Fd probe.

**Figure 2.**
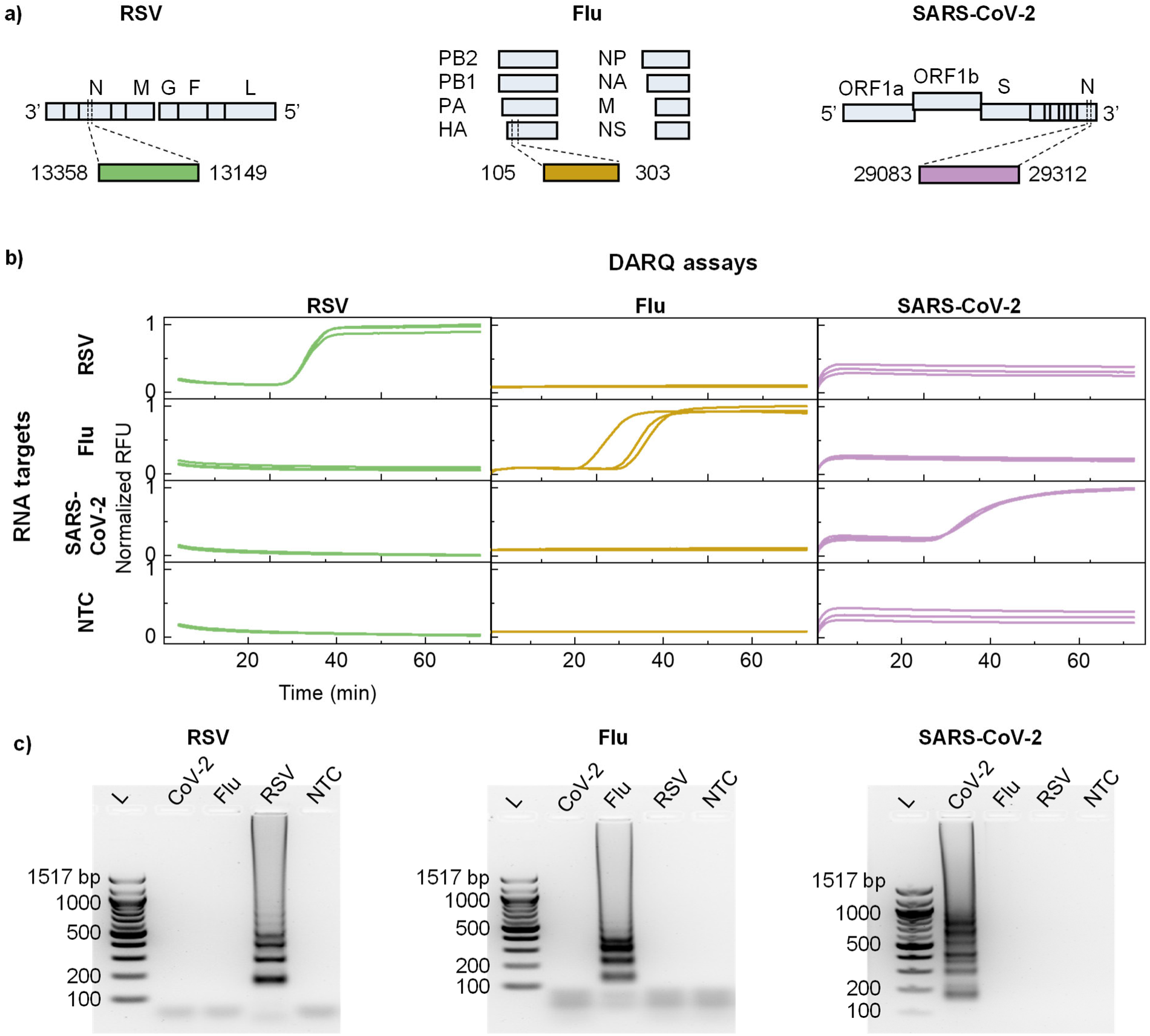
Preliminary validation of the DARQ RT-LAMP primer sets. (a) Specific regions of the three RNA targets that are amplified and detected during the single-plex DARQ RT-LAMP reactions, N, HA, and N1 for RSV, Flu, and SARS-CoV-2, respectively. Modifications were made to previously reported FIPs, incorporating a quencher at the 5’ end (QFIP) and adding a complementary fluorescence-labeled Fd probe to the reaction mix. (b) The normalized amplification curves (triplicates) to present the results of cross-reactivity tests for the DARQ RT-LAMP single-plex assays for 1000 copies/rxn of RSV, Flu, and SARS-CoV-2 RNAs as input targets separately and NTC on a benchtop thermal cycler. The matrix of amplification curves shows no cross-reactivity. (c) Images of gel electrophoresis performed on reaction products from subfigure b. The typical ladder patterns visible only for the intended targets in the case of all three assays corroborate the real-time results.

These primer and probe sequences are listed in **Supplementary Table 1**. A preliminary inspection of all the sequences using online tools such as NCBI BLAST, IDT’s Oligo Analyzer, and ThermoFischer’s Multiple Primer Analyzer did not indicate any non-target amplification and any significant probability for false amplification due to primer-primer interaction or primer-dimer formations.

To verify this experimentally, we conducted cross-reactivity tests for the single-plex assays on a benchtop thermal cycler. To test the cross-reactivity, we ran the DARQ RT-LAMP assays with 1000 copies/rxn of RSV, Flu, and SARS-CoV-2 RNAs as input targets separately. The normalized amplification curves (triplicates) for all combinations (three assays and four targets, each including water as NTC) are displayed as a matrix in **Figure 2b**. Amplification curves only along the diagonal of the matrix suggest only specific amplification of the intended RNA target for all assays without any false positives. This confirms the specificity of each assay, demonstrating no cross-reactivity with other target sequences, and serves as a fundamental validation of assay specificity. As a confirmatory test, we subjected all the reaction products to gel electrophoresis, and the results of all the assays are shown in **Figure 2c**. The typical ladder patterns visible only for the intended targets in the case of all three assays corroborate the real-time results. These results confirm that none of the assays show any cross-reactivity for the targets considered in this study and may be used further for a multiplexed detection assay.

### Performance of single-plex DARQ RT-LAMP assays on the analyzer

After confirming the absence of cross-reactivity in the DARQ assays against non-target RNAs, we evaluated the individual performance of each DARQ RT-LAMP assay by attempting to detect serially diluted target RNA on the developed analyzer and benchtop thermal cycler parallelly. We used target RNAs in the range 1.5x10^5^ to 15 copies per reaction (cp/rxn) with 10x dilution and then in the range 750 to 93.75 cp/rxn with 2x dilution in triplicate reactions for all concentrations in the RSV, Flu, and SARS-CoV-2 assays. The results for baseline subtracted amplification curves for the RSV assay obtained on the analyzer are presented in **Figure 3a**. The threshold line (black dotted) is calculated such that C_th_ = µ + 3σ where µ and σ are the mean and standard deviation of the predicted fluorophore concentration for the NTC reactions. This threshold concentration was calculated separately for each assay to measure the times to positive (T_p_) for all reactions that showed amplification. The mean times to positive as a function of RNA concentration are shown in **Figure 3b** with linear fit having an R^2^ value of 0.91.

**Figure 3.**
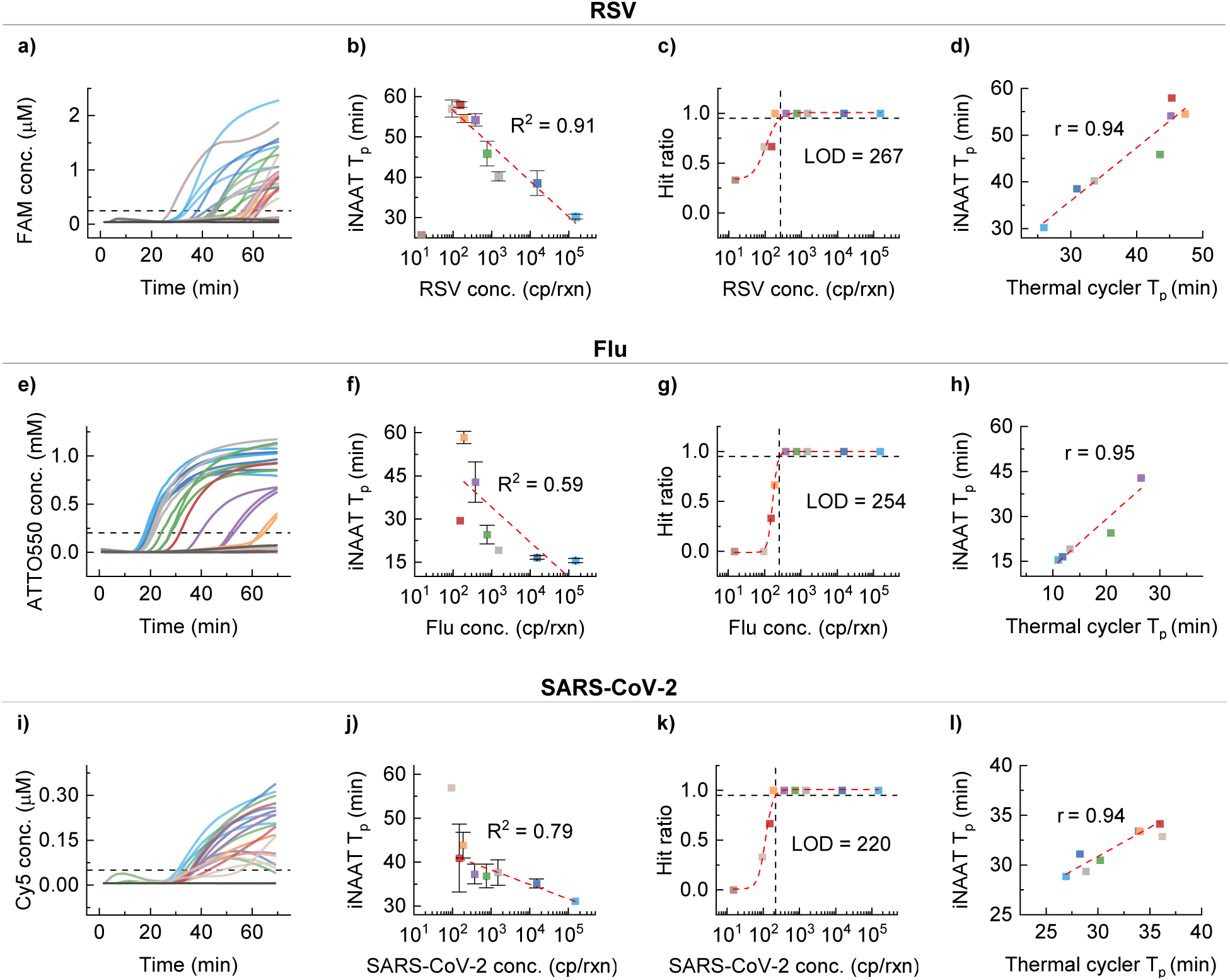
Analytical characteristics of the DARQ single-plex assays performed on the analyzer. (a) Amplification curves for the RSV assay, spanning a range of serially diluted RSV RNA concentrations, 10x serial dilution between 1.5x10^5^ and 15 copies/rxn, followed by 2x serial dilution between 750 and 93.75 copies/rxn. The horizontal dashed line represents the threshold concentration computed as µ+3σ, where µ is the mean, and σ is the standard deviation for NTC reactions. (b) Summary of the time to positive (T_p_) for triplicate reactions at each RNA concentration with a linear fit of R^2^ = 0.91. (c) The hit ratio for these serially diluted samples and a logistic fit reveal a limit of detection of 267 copies/rxn. (d) A scatter plot comparing the T_p_ seen on the analyzer with the T_p_ seen on a benchtop thermal cycler suggests good collinearity with Pearson’s r = 0.94. (e), (f), (g), and (h) present the same information for the Flu DARQ assay and reveal that the linear response between T_p_ and RNA concentration has R^2^ = 0.59 with a LOD of 254 copies/rxn and collinearity with thermal cycler has a Pearson’s r = 0.95. (i), (j), (k), and (l) present the performance of the SARS-CoV-2 assay that has a linear response between T_p_ and RNA concentration with R^2^ = 0.79 and LOD of 220 copies/rxn and collinearity with thermal cycler has a Pearson’s r = 0.94.

To estimate the LOD of the assay, we examined the hit ratios at various RNA concentrations, which are given as the number of positive tests over the total number of tests at a particular concentration and presented in **Figure 3c**. The experimental hit ratio data was fit with a logistic curve to reveal the LOD as 267 at the 95% confidence level when 1.5 µL RNA was used in a 25 µL reaction. This LOD is higher than some previously reported assays and is likely a result of poor reaction efficiency. However, considering a mean concentration of ∼6x10^3^ cp/µL of RSV(51) in a raw sample or ∼3000 cp/rxn (when extracted RNA is concentrated to 100 µL), it should be sufficient for qualitative detection in a multiplexed assay. **Figure 3d** compares the times to positive seen on the analyzer and a benchtop thermal cycler at various concentrations. A Pearson’s r value of 0.94 for RSV suggests that the analyzer’s operation is independent of the assay, and the efficiency and LOD observed are assay artifacts not affected by the analyzer. Please refer to **Supplementary Figure S2** for the amplification reactions performed on the benchtop thermal cycler.

Similarly, **Figures 3e-h** present the performance of the Flu assay with a linearity (R^2^) of 0.59, a LOD of 254 cp/rxn, and a Pearson’s r of 0.94 when benchmarking the analyzer against a benchtop thermal cycler. Considering a mean concentration of ∼3.1x10^3^ cp/µL for Flu (52) in clinical samples, it should be sufficient. **Figures 3i-l** present information for the SARS-CoV-2 assay, which has a linearity (R^2^) of 0.79, a LOD of 220 cp/rxn, and a Pearson’s r of 0.91, considering ∼10^5^ cp/µL for target RNA in SARS-CoV-2 samples(53), this LOD would be sufficient. Although these results indicate a semi-quantitative ability and demonstrate varying efficiencies for each assay, they show the performance of the single-plex assays, the analyzer’s ability to amplify target RNAs, and its comparable performance to a benchtop thermal cycler. Although a systematic comparison between the efficiencies of DARQ and standard Syto-9-based assays (keeping all other variables the same) was not attempted in this or prior studies, the involvement of a fluorophore-attached primer in the amplification pathway may adversely affect efficiency, linearity, and LOD. However, this finding is not expected to impact the qualitative detection capabilities of the subsequent multiplexed assay, which is the goal of this study.

### Effect of multiplexing DARQ assays on the detection of a single RNA target

After evaluating the single-plex assay performances, we set out to test the impact of multiplexing the primer sets on detecting a single RNA target of fixed concentration. We tried to amplify 750 cp/rxn of RSV, Flu, and SARS-CoV-2 purified RNAs independently using the single-plex, 2-plex, and 3-plex assays. To depict the outcomes of these experiments, the baseline subtracted amplification curves of eight replicates for the positive and negative control reactions in different primer set conditions are shown in **Figures 4a to 4c**.

**Figure 4.**
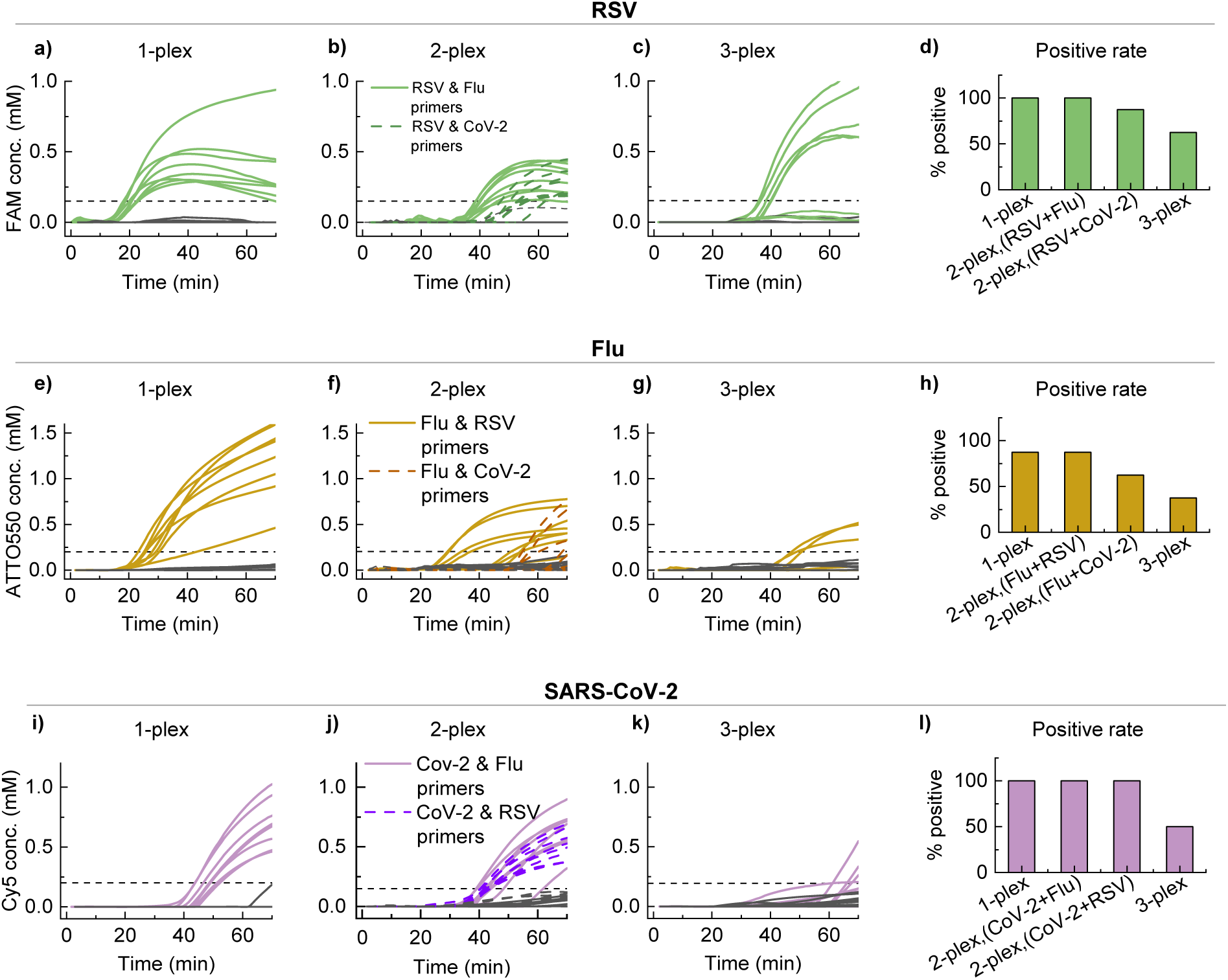
Evaluating the effect of increasing the number of distinct primer sets on the detection of a target RNA. (a), (b) and (c) Present the amplification curves of the 1- plex, 2-plex, and 3-plex assays when 750 copies/rxn of RSV RNA (green) and negative controls (dark gray) were added as input to the reaction in eight replicates. (d) The positive detection rate for all assays when targetting RSV. (e), (f) and (g) Present the amplification curves for the same set of primers as in a, b, and c when 750 copies/rxn of Flu RNA (yellow) and negative controls (dark gray) were added as input. (h) The positive detection rate for all assays when targetting Flu. (i), (j), and (k) Present the amplification curves for 750 copies/rxn of SARS-CoV-2 RNA (purple), and negative controls (dark gray) were added as input. (l) The positive detection rate for all assays when targetting SARS-CoV-2. In general, the detection rate decreases for all targets as the number of primer sets increases despite minimal assay optimization. This is likely due to a reduction in the concentration of the primers that target the specific RNA when the primer sets are multiplexed together to keep the total primer concentration commensurate with the amount of DNA polymerase.

When multiplexing primer sets together, certain assay parameters needed optimization to prevent false positives due to increased primer-primer interaction and to enable enzyme activity similar to the single-plex assay. For the 2-plex assay in **Figure 4b**, the primer concentrations were reduced to 50% of the single-plex assay concentrations **(Figure 4a)**. Similarly, for the 3-plex assay **(Figure 4c)**, the primer concentrations were reduced to 33% of the single-plex assay concentrations, or 1/n of the usual concentration, where n is the number of primers sets in the multiplexed assay. Another condition that was optimized was the ratio of FIP:QFIP, which was changed to ∼1.5:1 instead of 1:1. Thus, the minimal assay optimization helped preserve the detection ability for RSV targets as the degree of multiplexity increased up to three, albeit poor efficiency as evidenced by the reducing number of reactions containing target showing amplification. A summary presented in **Figure 4d** by plotting the percentage positive rate for the detection of RSV RNA in the presence of different primer sets shows that the detection rate decreases as more primer sets are added.

On the same lines, the detection rates of Flu (**Figures 4e to 4h**) and SARS-CoV-2 (**Figures 4i to 4l**) also decreased as the number of primer sets added to the assay increased, most likely due to the reduced primer concentrations. While slight differences in detection rates across targets exist, the 3-plex assay demonstrated clear detection capability for RSV, Flu, and SARS-CoV-2 replicates. These results show that although the sensitivity of detection for each target may be affected due to the reduced primer concentrations, no false positives due to primer-primer interaction were seen in any of the multiplexed assays, and the 3-plex assay may be used for detecting multiple targets in the same reaction.

### 3-plex assay to detect multiple RNA targets

In the previous section, we demonstrated the adverse effect of increasing the number of primer sets on detecting a single RNA target of fixed concentration due to the reduced concentrations of individual primers. This section evaluates the 3-plex assay’s ability to detect multiple RNA targets simultaneously, simulating a situation where a patient may have a co-infection of two or more respiratory pathogens. Since any fluorophore (attached to the specific probe) has an emission spectrum rather than emission at a single wavelength, multiple channels of the spectrophotometer may detect this emission, and the emission spectra from multiple fluorophores in a reaction tube may overlap, preventing an easy readout. To resolve these overlapped emission spectra, we developed a Multilayer Perceptron (MLP) Neural Network to predict the time-varying fluorophore concentration of the 3-plex RT-LAMP assay(48).

The 3-plex assay was used to detect 1500 cp/rxn of each RNA target in eight different combinations: individual targets, two targets, all three targets, and NTC. The raw fluorescence data acquired by the analyzer was fed to the NN to predict the time-varying fluorophore concentrations for the progressing reaction. The predicted fluorophore concentrations for eight replicates of each combination are presented as real-time amplification curves for RSV, Flu, and SARS-CoV-2 in **Figures 5a** to **5c**, respectively. Although no clear trend was seen among times to positive, even if multiple targets are present and accurately detected by the 3-plex assay, we notice certain false positives and negatives for each target. The assay’s detection ability is summarized in **Figure 5d** using a binary comparison plot, which compares the presence or absence of input RNA targets (RSV, Flu, and SARS-CoV-2) with the assay’s detection results. This figure visually represents whether the assay successfully detected the input target RNA for each sample, highlighting its ability to identify single or multiplexed targets. Further illustrating the assay’s performance, **Figure 5e** presents a confusion matrix that quantifies the number of correct target-wise classifications ranging between zero and eight across different RNA combinations. The overall classification accuracy of the assay is 80%, with the highest number of misclassifications seen for the combination that has all three RNAs present together. This may suggest variability in amplification kinetics across combinations and preferential amplification of certain targets not leaving enough resources for others. Thus, this figure demonstrates the ability of the 3-plex assay to detect all the RNA targets, whether present individually or with another target, and it can be used for multiplexed and co-infection detection.

**Figure 5.**
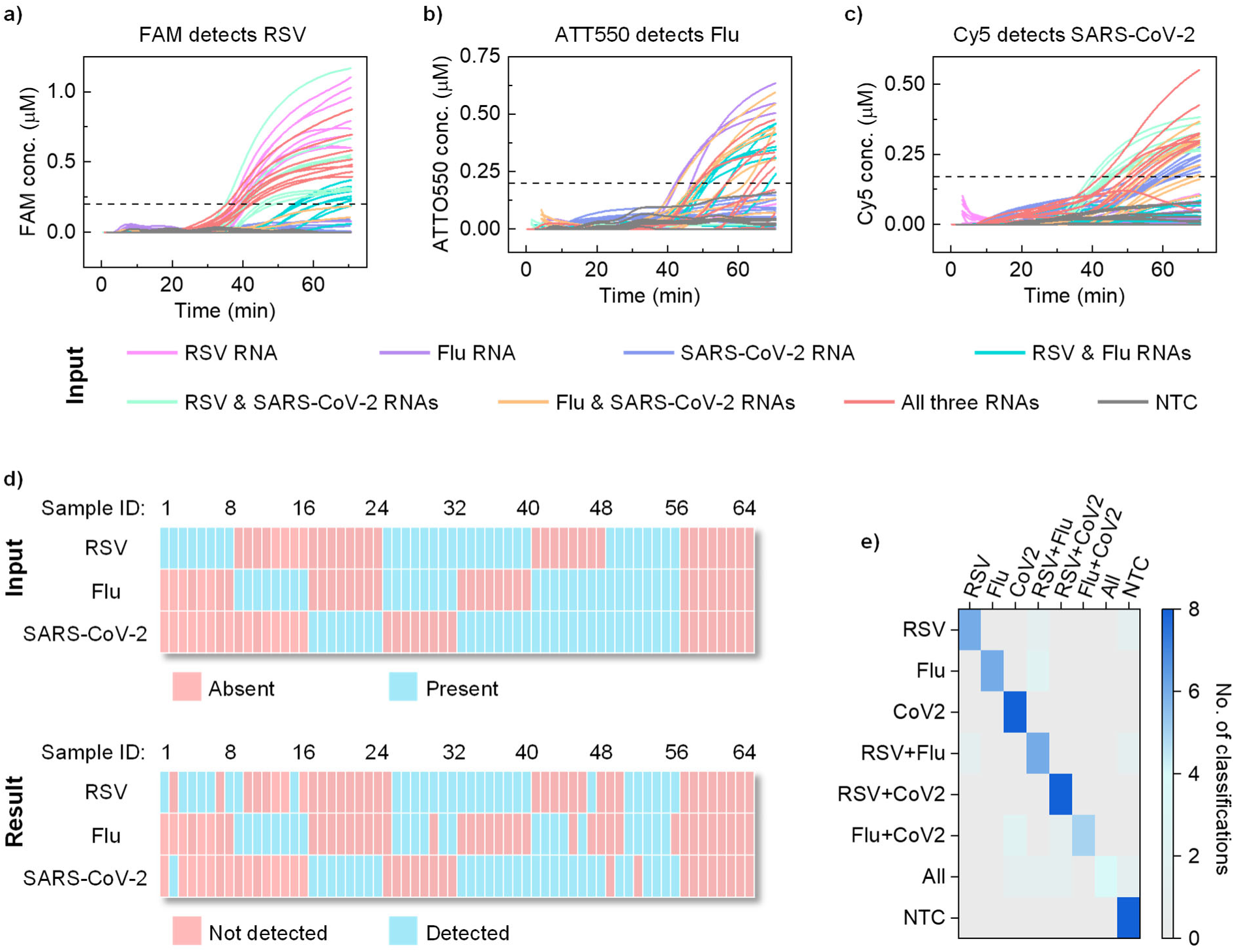
The ability of the 3-plex DARQ RT-LAMP assay to correctly classify single or combinatorial purified RNA input. (a) The amplification curves for detection of RSV RNA marked by the FAM fluorophore for various (eight) scenarios, including single RNA targets, dual RNA targets, all RNA targets (concentration: 1500 cp/rxn), and NTC. The horizontal dashed line represents the threshold concentration computed as µ+3σ, where µ is the mean, and σ is the standard deviation for NTC reactions. (b) The amplification curves for detection of Flu RNA marked by the ATTO550 fluorophore for the same reactions as in a. (c) The amplification curves for detection of SARS-CoV-2 RNA marked by the Cy5 fluorophore for the same reactions as in a. (d) Binary comparison plot between input RNA targets and assay detection results. The top panel represents the input matrix, showing the presence or absence of RSV, Flu, or SARS-CoV-2 RNA in each sample (eight replicates for eight scenarios). The bottom panel displays the detection results from the 3-plex DARQ RT-LAMP assay. Blue indicates presence/detection, while pink represents absence/non-detection. (e) Confusion matrix summarizing the assay’s classification accuracy, where each cell represents the number of classifications for a specific combination of input RNA targets (range: zero to eight). This demonstrates the assay’s ability to accurately detect single and multiplexed RNA targets, with an overall detection accuracy of 80%.

### Spiked saliva sample testing

After validating that the 3-plex assay can detect single and multiple targets using the respective purified RNAs, we attempted to detect single and co-infections in mock saliva samples. The mock samples were prepared by spiking purified RNA in negative saliva obtained from volunteers such that each target was present at a clinically relevant concentration of 10^5^ cps/mL of saliva. 200 µL of such spiked saliva was used to extract and purify the viral RNA by utilizing the Qiagen QIAamp viral RNA mini kit and a portable centrifuge to elute a final volume of 60 µL. In a separate work, we have demonstrated the use of this custom-developed portable centrifuge(54) with an ∼85% efficiency for use with a column-based extraction kit and a performance comparable to a benchtop centrifuge. Thus, we expect a viral RNA concentration of ∼1400 cp/µL of eluate from 200 µL of raw saliva and ∼6300 cp/rxn when 4.5 µL eluate is used in each 3-plex reaction.

After verifying satisfactory RNA extraction, the 14 mock samples having one, two, or no RNA targets were first tested using 1-plex RT-PCR (triplicates) as a gold standard. The amplification curves for RSV, Flu, and SARS-CoV-2, shown in **Supplementary Figure S3**, indicate accurate identification and quantification of the target RNA in each mock sample. Then, the same samples were tested using the 3-plex RT-LAMP assay in triplicates. **Figures 6a to 6c** show the predicted fluorophore concentrations representing the real-time amplification curves for each target of the 3-plex assay. Binary comparison plots in **Figure 6d** show that the 3-plex RT-LAMP assay classification matched with RT-PCR for 32 out of 42 tests (14 mock samples tested in triplicates), i.e., 76% agreement. This is further summarized by plotting the receiver operating characteristic (ROC) curves for each target separately, and the area under the curve values (AUC) values are 0.82, 0.93, and 0.96 for RSV, Influenza, and SARS-CoV-2, respectively. Thus, this section demonstrates the ability of the DARQ 3-plex assay to qualitatively identify single or co-infection of RSV, Flu, or SARS-CoV-2 targets in saliva samples by using a single reaction on the portable analyzer and fully ready to be implemented at the point-of-need. Please note we did not attempt the detection of three targets in saliva as it would be a highly improbable situation where a patient is infected with three different viruses at the same time; however, the previous section provides sufficient evidence that this 3-plex test would be able to identify it as well.

**Figure 6.**
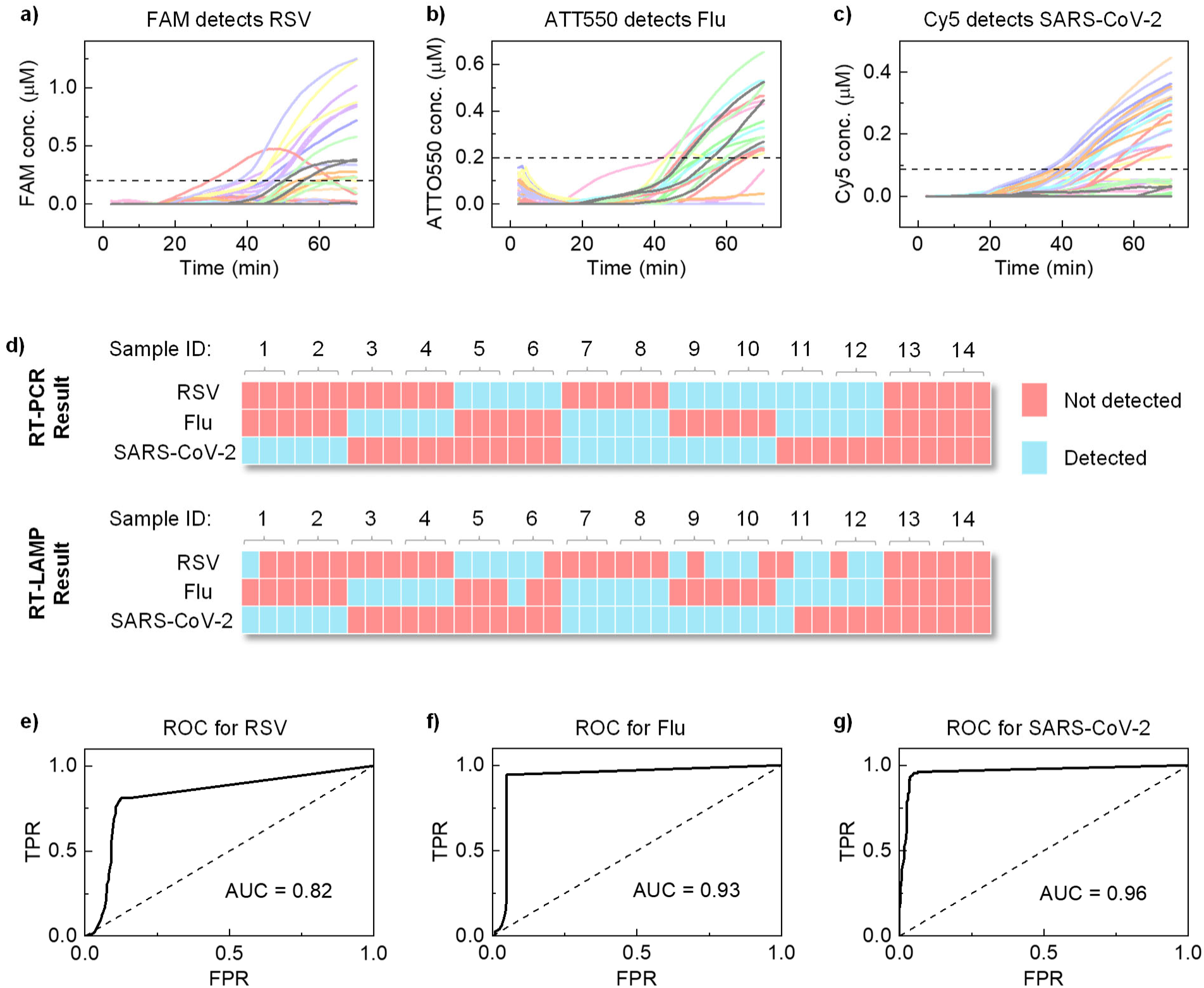
Mock saliva sample testing using the developed 3-plex RT-LAMP assay. Mock saliva samples were prepared by spiking purified RNA in negative saliva obtained from volunteers such that each target is present at a clinically relevant concentration of 10^5^ cp/mL of saliva. (a), (b) and (c) Present the real-time amplification curves for detecting RSV, Flu, and SARS-CoV-2 by the 3-plex DARQ RT-LAMP assay on the analyzer. (d) Binary comparison plot demonstrating 76 % agreement between the 3-plex RT-LAMP assay and RT-PCR when 14 mock samples were tested in triplicates. (e), (f) and (g) display the Receiver Operating Characteristic (ROC) curves for RSV, Flu, and SARS- CoV-2, respectively, comparing the performance of 3-plex RT-LAMP against 1-plex RT- PCR as the ground truth. TPR: True Positive Rate, FPR: False Positive Rate.

## Discussion

Point-of-need tests for respiratory symptoms should ideally be minimally invasive, provide rapid sample-to-answer results, and be easy to administer. This approach allows for quick differentiation between infections with similar symptoms while reducing patient hesitance and minimizing the risk of injury. A prompt diagnosis also enables timely, targeted treatment within optimal therapeutic windows, enhancing effectiveness. Conducting a panel of targets in such multiplexed tests on saliva allows for the simultaneous detection of multiple pathogens, including co-infections, further supporting the comprehensive and accurate diagnosis.

In this work, we developed a multiplexed diagnostic test designed to detect SARS-CoV-2, Flu, and RSV RNA from non-invasive saliva samples, directly addressing the challenge of distinguishing between respiratory viral infections with overlapping symptoms at the point of need. Our approach tackles the issue of differentiating overlapping spectral signals from multiple fluorophores within the same reaction tube by first gathering calibration data for each fluorophore and then training a neural network model to estimate time-varying fluorophore concentrations in a multiplexed RT-LAMP assay using a simple, low-cost device that does not rely on specialized optical filters, unlike other examples(30, 55). Future efforts would focus on refining the neural network model to leverage calibration data from individual fluorophores, which could enable the model to predict combined behaviors of fluorophores, reducing the need for extensive combinatorial experiments.

In any diagnostic test, it is essential to evaluate assay performance, particularly when using a multiplexed assay with multiple targets. Here, we demonstrated the use of a DARQ RT-LAMP assay in a portable analyzer for qualitative detection, as serial dilution tests of purified viral RNA suggest sub-optimal limits of detection, which are nonetheless sufficient for qualitative assessment. Previous studies have indicated that quencher- probe duplex (QPD) methods may inhibit amplification depending on QPD concentration relative to the supplemented primer(32, 45). Despite the optimization of QPD and other assay components, the LOD may not match that of the intercalating dye version of the assay(56). Although not thoroughly evaluated, this limitation may stem from the need to maintain adequate fluorescence detection, which requires a certain concentration of the quencher-probe duplex (QPD). Reducing the QPD concentration too much could compromise fluorescence detection, while the necessary concentration may, in turn, reduce amplification efficiency. To improve diagnostic sensitivity, especially in QPD- based approaches, enhancing primer design and exploring different quencher and probe placements may offer a pathway to achieve a better LOD.

The absence of false positives in our multiplex RT-LAMP assay demonstrates its analytical specificity, which is necessary when multiple primer sets are utilized. By effectively adjusting primer concentrations, we prevent false amplification due to primer-primer interactions; our approach enhances the reliability of results, which is essential for accurate pathogen screening. However, the observed reduction in positivity rate points to a sensitivity trade-off. Future research should aim to optimize the assay conditions, focusing on improving the detection limit. Enhancing LOD is crucial for applications requiring early detection, such as during the onset of an outbreak or in low-resource settings where laboratory equipment is limited.

Employing the multiplex RT-LAMP in a single-pot format for testing multiple RNA targets demonstrated the assay’s robustness and adaptability, which aligns with the emerging need for flexible diagnostic platforms capable of simultaneously detecting and distinguishing multiple pathogens. Our results show that while the assay can qualitatively detect viral RNAs, it aligns well with the performance of standard RT-PCR, confirming its practical utility. Another aspect not explored in this study is the quantitative analysis during multiplexing due to insufficient sensitivity, which could impede the detection of low viral loads. Future studies should, therefore, focus on investigating the kinetic impacts of multiplexing in RT-LAMP assays, such as how the presence and concentration of one target influence the detection of another. Additionally, it is essential to explore methods to enhance the quantitative capabilities of these assays, improving their ability to accurately measure viral loads and integrate minimal sample preparation steps, particularly for saliva(40), which could extend the assay’s applicability to more routine diagnostic use.

This study contributes to the ongoing development of molecular diagnostics by enhancing the flexibility and specificity of multiplex RT-LAMP assays. The ability to maintain specificity while testing for multiple pathogens in a single assay setup could position this approach as a practical tool in outbreak management and disease surveillance with simple-to-obtain samples that can improve compliance. Enhancements in assay sensitivity and integration of streamlined sample preparation methods will further the practical application of this technology. Our findings lay the groundwork for future innovations in point-of-need diagnostics, promising improved healthcare outcomes through early and precise pathogen detection.

## Materials and methods

### Analyzer design and fabrication

The analyzer housing, adapters to mount Printed Circuit Boards (PCBs), and the machined aluminum heating block were designed using SolidWorks CAD software. Structural parts were fabricated using MakerBot MethodX 3D printer (Brooklyn, NY) with MakerBot ABS (acrylonitrile butadiene styrene) material, and the heating block was machined by Protolabs Network. The heating block uses four two- ohm power resistors (MP725-2.00) mounted using a thermally conductive adhesive paste (Arctic Alumina) and an MC65F103A 10 k-ohm thermistor (Amphenol Thermometrics, St. Mary’s, PA) in a small recess for temperature feedback. PCBs were designed using AutoDesk Eagle CAD software and fabricated by OSH Park LLC (Lake Oswego, OR). The excitation LEDs (SK6812) were purchased from Adafruit Industries (New York, NY), whereas other electronic components such as the Raspberry Pi Zero, power resistors, thermistor, and spectral sensors (AS7341) were purchased from DigiKey.com. A cost estimate of the analyzer is provided in **Supplementary Table S3.**

### Target RNAs

Heat-inactivated SARS-CoV-2 (ATCC VR-1986HK) RNA and Quantitative Genomic RNAs of Influenza A virus (H1N1) strain A/PR/8/34 (ATCC VR- 95DQ) and Human Respiratory Syncytial Virus strain A2 (ATCC VR-1540DQ) were purchased from American Type Culture Collection (Manassas, VA).

### RT-LAMP assay

The RT-LAMP reaction mix consists of 1x isothermal buffer (20 mM Tris-HCl, 10 mM (NH_4_)_2_SO_4_, 50 mM KCl, 2 mM MgSO_4_, 0.1% Tween 20, pH 8.8), 1-plex, 2-plex or 3-plex primers, 0.5 M Betain, 6 mM MgSO_4_, 1.4 mM deoxyribonucleotide triphosphates (dNTPs), 0.5 U/µL Bst 2.0 DNA polymerase, 0.3 U/µL WarmStart Reverse Transcriptase, 1.5 µL purified RNA template (per target for multiplexed detection), 6 µM syto-9 (for applicable assay) and PCR grade water to bring total reaction volume to 25 µL. The LAMP assay was performed at a constant temperature of ∼62°C. The 1-plex RT-LAMP assay using syto-9 intercalating fluorophore consists of the usual primer concentrations, 0.2 µM F3 and B3, 0.8, LF and LB, and 1.6 µM FIP and BIP; however, the DARQ assay substitutes 1.6 µM FIP and 6 µM syto-9 with 0.8 µM FIP and QFIP, and 0.65 µM Fd. For the multiplexed assays, the primer concentrations for all 6 primers for each target were reduced by 1/n, such that n represents the number of primer sets or targets. Thus, for the 2-plex assay, the primer concentrations are 0.1 µM F3 and B3, 0.4 µM FIP, QFIP, LF, and LB, 0.325 µM Fd, and 0.8 µM BIP. The tri-plex assay, however, required optimizing the ratio of FIP:QFIP, such that FIP is in excess, resulting in the following primer concentrations, 0.067 µM F3 and B3, 0.4 µM FIP, 0.267 µM QFIP, LF and LB, 0.21 µM Fd and 0.53 µM BIP.

### RT-PCR assay

The 20 μL RT-PCR reaction consists of 10 µL of qScript XLT One-Step RT-qPCR Tough Mix (2X), 1.5 µL of premixed primers, and a probe for detecting SARS-CoV-2 or 0.4 µM forward and reverse primers and 0.1 µM probe for detecting Flu and RSV, 3.5 µl of the RNA template, and 3.5 µl of H_2_O. 2019-nCoV RUO Kit, catalog #10006713, for detecting SARS-CoV-2 and custom-designed primers and probes for Flu and RSV, detailed in **Supplementary Table S2**, were purchased from Integrated DNA Technologies, Coralville, USA. The RT-PCR involved cDNA synthesis at 50 °C for 10 min, initial denaturation at 95 °C for 1 min, followed by 45 cycles of a 5-second denaturation step at 95 °C and a 30-second annealing step at 55 °C in a Bio-Rad CFX96 Real-Time PCR system.

### Data processing and analysis

The predicted fluorophore concentrations (*C*) were processed using baseline subtraction, similar to the method used by the CFX 96 benchtop thermal cycler. A linear fit was calculated for the values between 5 and 10 min, and the slope (*m*) and intercept (*b*) values were used to determine the subtraction value (*SV*): *SV=t×m+b* for any time *(t)*. The baseline-subtracted concentrations (*C_bs_*) were then computed as *C_bs_*=*C*−*SV* and smoothed using an Exponential Moving Average (EMA) filter implemented in Python using the pandas library, with any negative values being set to zero to ensure non-negative fluorescence readings.

## Acknowledgments

This work was partially supported by the National Institutes of Health (R61AI147419, R33AI147419, R33HD105610), the National Science Foundation (2319913), and the American Rescue Plan Act through USDA APHIS (APHIS/NIFA Collaborative award# 2023-70432-41395). Any opinions, findings, conclusions, or recommendations expressed in this work are those of the authors and should not be construed to represent any official NSF, NIH, USDA, or US Government determination or policy.

